# Compartment size and proliferation shape the murine T cell receptor repertoire in splenic T- and B cell zones during an immune response

**DOI:** 10.1101/2025.03.28.645898

**Authors:** Franka Buytenhuijs, Cornelia Tune, Andrea Schampel, Lisa-Kristin Schierloh, René Pagel, Jürgen Westermann, Johannes Textor

## Abstract

The adaptive immune system recognizes billions of unique antigens using highly variable T cell receptors (TCRs). To what extent TCR features shape the spatial distribution of the TCR repertoire (TCR-R) within lymphoid organs remains unclear. Here, we compared the TCR-R composition in T cell zones (TCZ), B cell zones (BCZ), and germinal centers (GC) of the spleen by analyzing the complementarity-determining region 3 of the beta chain (CDR3β) in microdissected splenic compartments from naïve and immunized mice.

While we found substantial differences in TCR-R composition between these compartments, these differences largely disappeared once the effects of compartment size and proliferation were accounted for: During steady state, the few T cells found in BCZs strongly resemble random subsets of those found in TCZs. After immunization, expanded clones started appearing in the TCZ at day 3 and later shifted to the BCZ. GCs harbored even further expanded clones that were grouped into clusters with high sequence similarity. TCR-R differences during the immune response were well explained by a Galton-Watson model of T cell proliferation, suggesting that this could be the central factor shaping the observed TCR-R differences. Similarly, while TCR length and closeness to germline appeared to differ between the compartments, our models of random distribution and proliferation explained these differences almost entirely.

Hence, our findings suggest that splenic TCR-Rs are shaped predominantly by compartment size and proliferation rates rather than TCR sequence features. This would imply that quantitative assessment of TCR-R dynamics across heterogeneous spatial compartments during an immune response is only possible when these effects are explicitly accounted for.

## Introduction

The adaptive immune system can recognize billions of unique antigens. This process relies on the T cell receptor (TCR) repertoire (TCR-R)–a diverse pool of TCRs expressed on the surface of T cells. The TCR, which is unique for each T cell clone, recognizes antigens presented by major histocompatibility complex (MHC) molecules, leading to T cell and subsequently B cell activation. Upon antigen recognition, individual T cell clones undergo clonal expansion and differentiation, which changes the composition of the TCR-R. The potential number TCRs that could be generated by somatic recombination is estimated to range from 10^15^ and 10^20^ distinct sequences,^1, 2^ yet the actual TCR-R per individual likely only includes on the order of 10^7^ different TCR sequences.^3^ Apart from physical constraints (10^16^ T cells would weigh as much as an elephant), diversity is also reduced due to stringent selection processes in the thymus, which shape the functional repertoire of mature T cells.^4, 5^ A large fraction of this TCR diversity is determined by the complementarity-determining region 3 of the TCRβ chain (CDR3β), which is the most variable region of the TCR heterodimer. The CDR3β is formed by random recombination of variable (V), diversity (D), and joining (J) gene segments during T cell development.^1,6^

Immune responses are initiated and coordinated within secondary lymphoid organs (SLO) such as lymph nodes and spleen. These organs serve as gathering places for immune cells, facilitating interactions between different cell types and the generation of effective immune responses.^7,8^ A key effector mechanism of adaptive immunity is the T cell-dependent B cell response, which plays a crucial role in the production of antibodies targeting diverse antigens. This response relies on the complex and tightly regulated interplay between T cells and B cells. Here, T follicular helper (Tfh) cells – a subset of CD4^+^ T cells – are the key players.^9–12^ Upon activation in the T cell zone (TCZ) they migrate towards the B cell zone (BCZ) – guided by their chemokine receptor CXCR5 – where they establish contact with B cells that have captured the same antigen.^13,14^ They provide activation signals which drive B cell differentiation. These activated B cells then form germinal centers (GCs) within the BCZs to produce high-affinity antibodies through somatic hypermutation.

Despite extensive research about the interactions between T- and B cells in SLOs, relatively little is known about the spatial organization and compartmentalization of TCR-Rs *within* SLOs, particularly in the BCZ and GC. Given the essential role of T cells in shaping immune responses, understanding the distribution, selection, and expansion of antigen-specific T cell clones in these compartments is crucial.^15^ To characterize and compare the TCR-Rs in different splenic compartments, we isolated the different compartments by laser-microdissection of murine splenic cryosections and performed next-generation sequencing (NGS) of the CDR3β region to separately characterize the TCR-Rs of each compartment in naïve state as well as during an immune response to sheep red blood cells (SRBC).^16–19^

The analysis of TCR-Rs presents a unique challenge due to their immense diversity and the significant variability in T cell numbers between different compartments. In particular, T cell densities are expected to differ by orders of magnitude between the TCZ, BCZ, and GC, which may influence the interpretation of repertoire dynamics. To account for the effects of these differences in population size during an the immune response, we applied a combination of downsampling and proliferation modeling. This approach allowed us to distinguish TCR-specific repertoire differences from changes induced by clonal expansion and compartmental redistribution. Our findings demonstrate that the observed differences between compartments can largely be attributed to variations in compartment size and distinct proliferation dynamics at different time points following an immune response.

## Methods and Materials

### Mouse model and experimental groups

All experiments were conducted in accordance with the German Animal Welfare Act and approved by the Animal Research Ethics Board of the respective Ministry of Environment (Kiel, Germany, no. 72-5/15 and 25-3/18; Stuttgart, Germany, no. M11/14). Eight- to twelve-week-old C57BL/6J mice were either immunized by injecting 200 μl phosphate-buffered saline (PBS) containing 10^9^ SRBC into the tail vein or 200 μl PBS as control. Animals were assigned randomly to the experimental groups. Control mice were sacrificed 3 days post-immunization (3d p.i.), and immunized mice were sacrificed 3d, 4d, and 7d p.i., respectively, resulting in four groups: ‘PBS’ (n=8), ‘SRBC 3d’ (n=8),‘SRBC 4d’ (n=8), and ‘SRBC 7d’ (n=6). Since we abstained from using naïve control groups also for the 4d and 7d time points, the PBS group was used as an overall control for all time points.

### Analysis and sequencing of cells in splenic cryosections

#### Ki67/B220 staining

Splenic cryosections were stained for proliferating cells and B cells using the markers Ki67 and B220, respectively. Sections were fixed using chloroform and acetone washed in TBS-Tween, followed by fixation for 45 minutes in 4% PFA at 4°C. Subsequently, the primary anti-mouse Ki67 antibody was applied overnight and detected with the secondary biotinylated anti-rat IgG antibody. ExtrAvidin Alkaline Phosphatase was added for 30 minutes, followed by a further washing step. To visualize proliferating cells, we applied Fast Red staining solution. Next, the primary anti-mouse B220 antibody was added for one hour and again detected with anti-rat IgG antibody and in a next step ExtrAvidin Alkaline Phosphatase. To visualize B lymphocytes, we used Fast Blue staining. Slides were covered with Aquatex. Digital images were taken using Axiophot Microscope and AxioCam (Carl Zeiss, Jena, Germany). Analyses were performed with ImageJ (National Institutes of Health).

#### TCRβ/B220 staining

As a control staining and to assess the number of T cells in different splenic compartments, cryosections on glass slides were stained for T and B lymphocytes using the markers TCRβ and B220, respectively. First cryosections were fixed using a methanol-acetone solution (1:1) for ten minutes at −20 °C. After purging for ten minutes in TBS-Tween, the primary biotinylated anti-mouse TCRβ antibody was applied for one hour. Unbound antibodies were washed away by TBS-Tween (ten minutes at room temperature) before the samples were incubated with ExtrAvidin Peroxidase for 30 minutes. After purging in TBS-Tween for 10 minutes again, DAB substrate was added for five minutes to visualize T cells. After the next washing step, the primary anti-mouse B220 antibody was added for one hour and remaining substances were washed away afterwards. The secondary biotinylated anti-rat IgG antibody was applied for 30 minutes then. Following purging in TBS-Tween for 20 minutes, the samples were incubated with ExtrAvidin Alkaline Phosphatase for 30 minutes and washed again.

To visualize B lymphocytes, we used Fast Blue staining solution for 25 minutes. After the last washing step, the slides were finally covered using Aquatex and cover slips and stored in the dark at room temperature. Examination of the splenic sections stained for TCRβ and B220 was performed using the Axiophot light microscope. The ImageJ software was then used to quantify the number of T lymphocytes present in the different splenic compartments. A calibration slide was used to adjust the right magnification by means of the function “Set scale”. Using the functions “Rectangle” and “Measure”, selections with a known area were transferred in TCZ, BCZ and GC regions. Within these sectors, TCRβ positive cells were counted separately, and number of T cells denoted in relation to an area of 1mm^2^ after averaging over several representative compartments per animal.

#### Laser microdissection

To analyze the different splenic compartments the Zeiss Axiovert 200M PALM Microbeam System was used to precisely isolate TCZ, BCZ and GC from cryosections stained with toluidine blue. Compartments were marked using PALM Robo Software (version 4.8) and then dissected by a pulsed UV laser. Isolated tissues were sampled in tubes whose lids were wetted with inert mineral oil. Subsequently, lysis buffer was added. For deep sequencing, separated compartments were dissected and 200 μl Lysis solution RL (Analytik Jena) per tube was added. An isolated area of at least 1 million μm^2^ per compartment was required to enable further processing, meaning that up to three slides per animal (covering 15 stained serial sections each) were used for dissection and sampled in individual tubes. After incubation for 30 minutes at room temperature the samples were stored at −20 °C.

#### RNA isolation and quantification

Total RNA of the isolated splenic compartments was extracted with the innuPREP RNAmini kit (Analytik Jena) and treated with DNase I (Sigma-Aldrich). RNA quantity was determined using the Quantus fluorometer (Promega Biosystems, Sunnyvale, CA).

#### CDR3 sequence analysis of the TCRβ-chain

Total RNA was extracted as described above and TCRβ-chain transcripts were amplified independently in a two-step reaction according to the manufacturer’s protocol (iRepertoire; patent no. 7,999,092). Gene-specific primers targeting each of the V and J genes were used for reverse transcription and first-round PCR (OneStep RT-PCRMix; Qiagen). In addition to a nested set of gene-specific primers, sequencing adaptors A and B for Illumina paired-end sequencing were added during second-round PCR (Multiplex PCR Kit; Qiagen). PCR products were run on a 2% agarose gel and purified using QIAquick Gel Extraction Kit (Qiagen). The obtained TCRβ libraries were quantified using the PerfeCTa-NGS-Quantification Kit according to manufacturer’s protocol (Quantabio) and sequenced using the Illumina MiSeq Reagent Kit v2 300-cycle (150 paired-end read; Illumina), gaining an average of ≈1.8 × 10^6^ reads per sample. The number of unique amino acid sequences ranged from approximately 50,000 to 110,000. Throughout this paper, different nucleotide sequence encodings for identical amino acid sequences are treated as equal and termed ‘clonotype’.

#### TCR repertoire generation

CDR3β identification, clonotype clustering and correction of sequencing errors were performed using MiTCR software. All parameters were set to the standard values of the ClonoCalc graphical user interface for MiTCR.^20^

#### Immunofluorescence staining to identify proliferating T cells

To assess the number of proliferating T cells within the different splenic compartments, cryosections were stained for proliferating cells (monoclonal αKi67; Alexa Fluor 594 rat anti-mouse/human Ki67 monoclonal antibody (11F6) BioLegend, USA) and CD4^+^ T cells (monoclonal αCD4; Alexa Fluor 488 rat anti-mouse CD4 monoclonal antibody (GK1.5) BioLegend, USA) in a humid chamber. Cryosections were fixed using 4% PFA in PBS and permeabilized in 0.5% Triton-X-100 in PBS. Blocking was performed in blocking solution containing 5% BSA and 5% normal mouse serum in PBS for one hour at room temperature. Both antibodies were applied at a concentration of 1:25 in blocking solution at 8 °C overnight. Sections were counterstained with Hoechst Dye (1:10,000 in PBS) and cover slipped with Moviol. For examination of the splenic sections stained for Ki67 and CD4 digital images were taken using a Keyence microscope. Analyses were performed with the ImageJ software (National Institutes of Health). A reference image was used to adjust the right magnification by means of the function “Set scale”. Using the functions “Rectangle” and “Measure”, selections with a known area were transferred in TCZ, BCZ and GC regions. Within these sectors, Ki67 and CD4 positive cells were counted separately. The number of positive cells was denoted in relation to an area of one mm^2^ after averaging over several representative compartments per animal.

### TCR-R data analysis

#### Downsampling

To ensure a fair comparison between the TCZ, BCZ, and GC, we downsampled the larger compartment by matching the number of *unique* amino acid sequences to that of the smaller compartment. This approach was only feasible when the TCZ was larger than the BCZ, which is why one mouse from the 4 day time point was excluded from the analysis. CDR3β amino acid sequences were sampled in batches of 50, with replacement. This iterative sampling continued until the number of unique amino acid sequences in the TCZ sample matched that in the BCZ, within a tolerance of ±25 sequences. The probability of selecting each sequence was weighted by the base-2 logarithm (log_2_) of its copy number, ensuring that more abundant sequences were more likely to be included in the sample. The final dataset represents a downsampled version of the TCZ (dsT) data for each mouse. Due to the stochastic nature of the process, the downsampling was repeated 10 times, and the results were averaged. The same downsampling procedure was applied to the BCZ samples to compare them with the GC data.

### Cell expansion modelling

Proliferation was modeled using a Galton-Watson branching process, a stochastic model widely used to describe population dynamics, particularly in terms of growth, survival, and extinction.^21^ In this model, each individual in the population, in this case, each clonotype, has a specific probability of producing offspring in each generation.

The process begins with the initial downsampled dataset in the first evolution step. During this step, each clonotype in the dataset attempts to proliferate, succeeding with a probability *x*. When proliferation is successful, the clonotype divides into two daughter cells. In subsequent evolution steps, the newly generated daughter cells from the previous step undergo the same proliferation process. This iterative cycle continues until no new cells are produced in a given evolution step, marking the conclusion of the proliferation cycle.

Once the cycle is completed, the total number of clonotypes generated is used to calculate a new copy number. In the case of positive expansion probabilities (*x*>0), the original copy numbers for each sequence are multiplied by the number of offspring produced, modeling a situation where proliferation is higher in the smaller compartment. For negative expansion (*x*<0), the original copy numbers are divided by the number of offspring produced. The negative probability models a situation where the expansion rate is higher in the larger compartment than in the smaller compartment.

### Statistical analysis

Because they have the highest risk of containing sequencing errors, sequences with only one mapped read were removed from all analyses. The distributions of unique clonotypes are compared using Kullback-Leibler (KL) divergence. The KL divergence is a statistical measure that quantifies the distance between two distributions, where zero means two distributions are equal. In the calculations of the KL divergences for the GC, the first 3 log_2_ copy numbers were not considered. Since our simplified proliferation model assumes that copy numbers can only increase when *x*>0, it cannot fit or accurately represent these low copy number values.

P values were computed using paired or unpaired t tests, as stated in the figure legends. All statistical analyses were performed within the R platform for statistical computing version 4.3.1. Sequence logos were created with R package ggseqlogo.^22^

## Results

### Similar CDR3β TCR repertoires in the T cell zone (TCZ) and B cell zone (BCZ) in the absence of antigen

To identify the TCZ and BCZ in murine spleens for microdissection and estimate the number of T cells per compartment, we used B220 and TCRβ staining. Initially, we stained spleen slices from eight naïve mice injected with PBS (**Figure 1A**). The density of T cells in the BCZ was ∼15-fold lower than in the TCZ: 1,072 ± 265 T cells/mm^2^ in the BCZ versus 16,370 ± 1,727 T cells/mm^2^ in the TCZ (**Figure 1B**). Using NGS of the CDR3β-chain, we assessed the number of distinct clonotypes in each compartment. The TCZ contained ∼55,000 unique clonotypes (54,055 ± 10,298), while the BCZ had ∼15,000 clonotypes (16,117 ± 6,330), corresponding to a 3.5-fold reduction in clonotype diversity in the BCZ compared to the TCZ (**Figure 1C**). Consequently, the T cells located in the naïve BCZ generate a distinct TCR-R that, although less diverse in absolute terms, is relatively more diverse and less redundant than the considerably larger TCR-R found in the TCZ.

**Figure 1.**
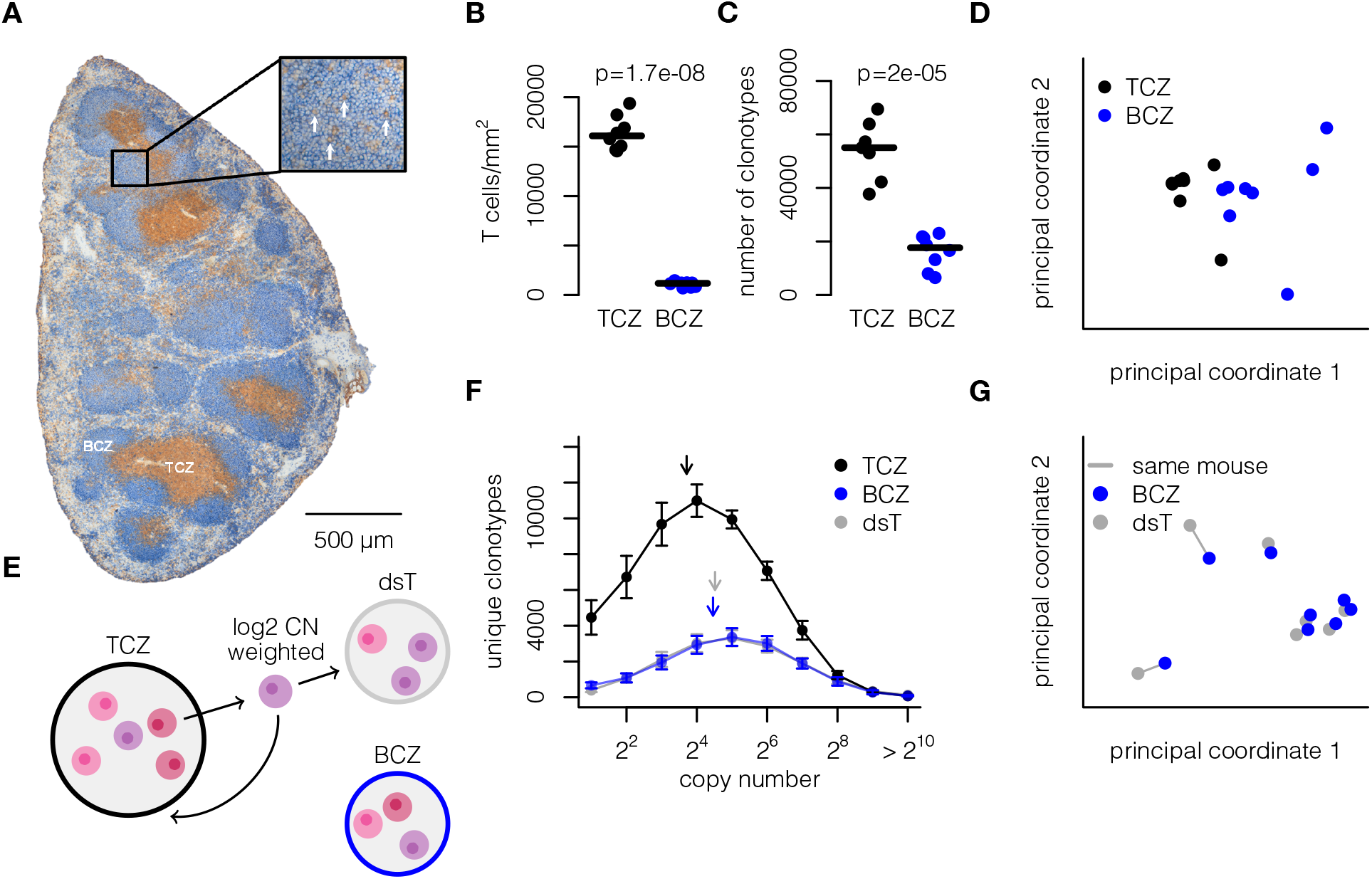
TCR-R differences between the T cell zone (TCZ) and B cell zone (BCZ) are largely attributable to the larger size of the TCZ. **(A)** Splenic section of a naïve mouse stained for T cells (TCRβ, brown) and B cells (B220, blue) to identify the TCZ and BCZ. The inset highlights T cells (TCRβ, brown, arrows) within the BCZ. **(B)** Number of T cells per section area (mm^2^) in the TCZ (black) and BCZ (blue), with means and individual animal data indicated (n=8 mice). P value: two-tailed paired t test. **(C)** Laser microdissection was used to isolate both TCZ (black) and BCZ (blue), followed by next-generation sequencing of the T cell receptor beta-chain to determine the number of different clonotypes (unique amino acid sequences); means and individual animal data are indicated (n=8 mice). P value: two-tailed paired t test. **(D)** Multidimensional scaling (MDS) overview of the TCZ and BCZ based on the overlap (Jaccard index) in junction amino acid sequence, V region, and J region. **(E)** For each mouse, a sample of unique amino acid CDR3β sequences from the TCZ was taken, weighted by the logarithm of the copy number, to match the number present in the BCZ. **(F)** Clonotype numbers within the TCZ (black) and BCZ (blue) are shown according to their copy number. dsT (grey) represents results for a TCZ downsampled to BCZ size. Geometric means are indicated by arrows (n=8 mice; average of 10 samples). **(G)** MDS overview comparing dsT and BCZ based on the overlap (Jaccard index) in junction amino acid sequence, V region, and J region.

To further investigate the differences between TCZ and BCZ repertoires, we assessed the similarity of samples within the compartments using multidimensional scaling (MDS) based on the Jaccard distance (1-Jaccard index), considering two clones to overlap if they shared the same junction amino acid sequence, V gene, and J gene usage (**Figure 1D**). This approach suggested that there were indeed some systematic differences between TCR-Rs from the BCZ and TCZ compartments. To determine to what extent these apparent differences were merely driven by the different numbers of T cells in the samples, we used a subsampling approach where we randomly selected the same number of clonotypes found in a BCZ sample from the TCZ sample for each animal (see Methods, **Figure 1E**).

When comparing the unique clonotype distributions across TCZ, BCZ, and downsampled TCZ (dsT),^23^ we found that the distribution of dsT closely resembled that of BCZ (**Figure 1F**). Additionally, when we conducted MDS analysis on samples from BCZ and dsT, we observed that samples from the same mouse clustered together (**Figure 1G**), eliminating the separation by compartment observed in the original comparison between BCZ and TCZ. These findings indicate that the observed differences between the TCZ and BCZ TCR-Rs are largely attributable to the larger size of the TCZ rather than inherent differences in TCR-R composition. Hence, in steady state, BCZ and TCZ have a very similar TCR-R clone size distribution once the difference in compartment size is taken into account.

### The differences between the TCZ and BCZ during an immune response are consistent with increased proliferation in the BCZ

Next, we examined the differences between the TCZ and BCZ at various time points after an immune response against SRBC using the same downsampling approach. At 3 and 4 days post-immunization (p.i.), the dsT did not align with the BCZ (**Figure 2A**), indicating that compartment size alone no longer explains the difference in clone size distribution during an immune response. We hypothesized that these discrepancies resulted from different expansion rates in the TCZ and BCZ. To model the effects of proliferation on the clone size distribution, we employed a Galton-Watson process,^21^ where each T cell could divide into two daughter cells at each evolutionary step (see Methods). At the end of the simulation, the copy numbers were adjusted according to the final count of generated daughter cells (see **Figure 2B**).

**Figure 2.**
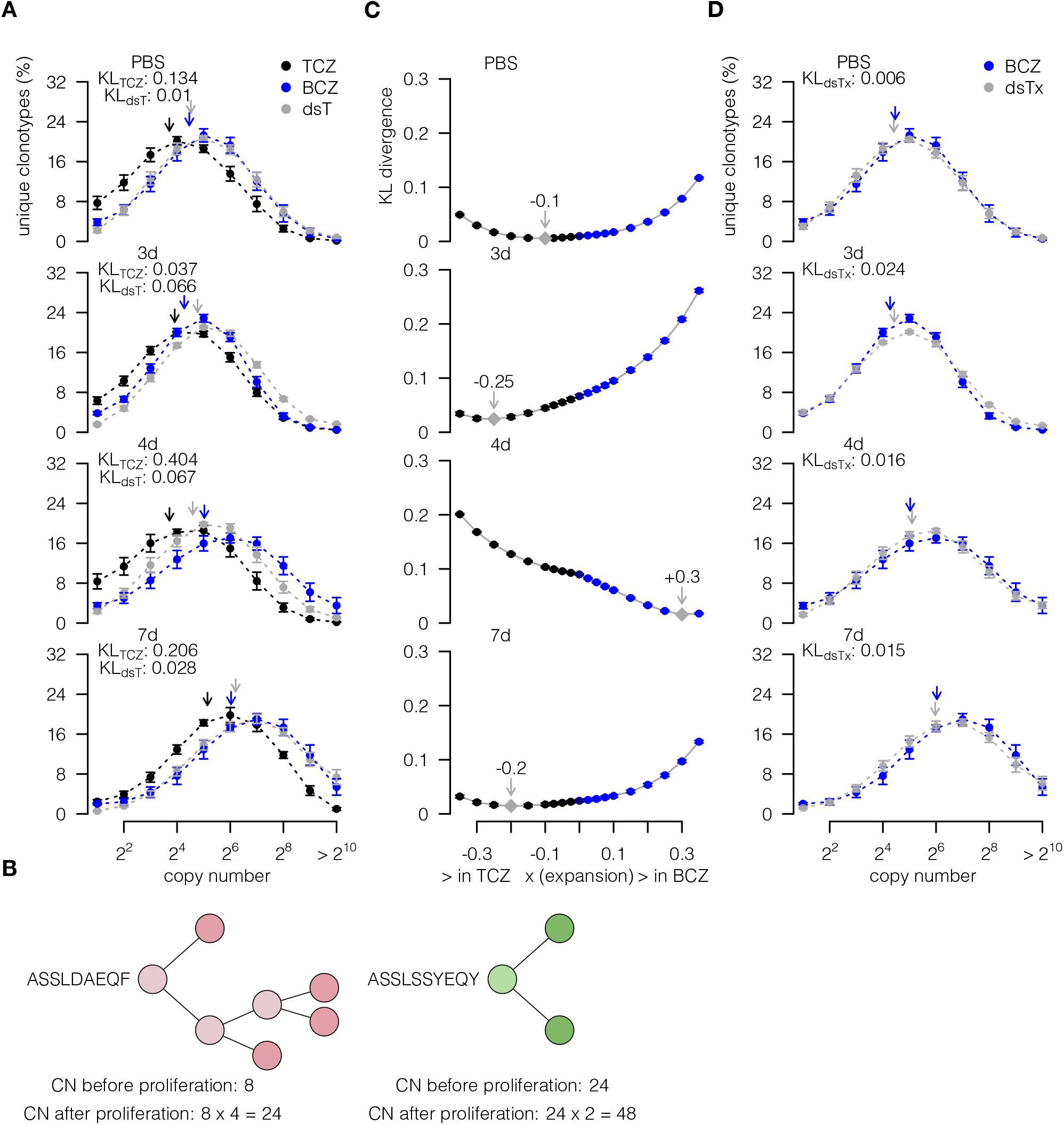
The differences between the T cell zone (TCZ) and B cell zone (BCZ) during an SRBC-induced immune response can be explained by varying expansion rates. **(A)** Clonotoype size distributions for PBS (n=8 mice), and at 3 days (n=6 mice), 4 days (n=7 mice), 7 days (n=6 mice), and 10 days (n=4 mice) post-immunization (p.i.). dsT refers to a TCZ downsampled to BCZ size. Geometric means are shown by arrows, bars depict standard errors. **(B)** Illustration of proliferation simulations using the Galton-Watson process. CN: copy number. **(C)** Kullback-Leibler (KL) divergence for different proliferation rates in TCZ samples at 3, 4, and 7 days p.i. with SRBC. Average of 10 samples. **(D)** Clonotype size distributions within the BCZ for PBS (n=8 mice), and at 3 (n=6 mice), 4 (n=7 mice), and 7 (n=6 mice) days p.i. dsTx (grey) shows the results for a downsampled TCZ with simulated expansion and the best fit (lowest KL) to the BCZ. Arrows indicate geometric means, error bars indicate standard errors.

To infer the proliferation rate from the clonotype distribution, we computed the KL divergence between the BCZ and dsTx for a range of different rates. At 3 days p.i., the best fit was obtained when applying a negative expansion rate (x = −0.25, KL = 0.014), indicating higher expansion in the TCZ than in the BCZ. In contrast, at 4 days p.i., the best fit was achieved with a high positive proliferation rate (x = 0.3, KL = 0.021), suggesting increased expansion in the BCZ, or extensive immigration of expanded clones from the TCZ. By 7 days p.i., the situation had largely reverted to that observed in the naïve state in that downsampling alone explained the BCZ data well again (PBS; **Figure 2C**). Using the best-fitting proliferation rates, the dsTx distribution closely resembled that of the BCZ (**Figure 2D**).

These findings suggest that the differences between the TCZ and BCZ clone size distributions during an immune response are not purely due to the difference in compartment size, but can be well explained by assuming early expansion of T cells in the TCZ, followed by a shift of expanded or proliferating clones to the BCZ at 4 days p.i.

### Germinal centers (GCs) harbor a distinct T cell receptor repertoire (TCR-R) characterized by high expansion rates

During an immune response, GCs emerge within the BCZ as specialized sites for B cell clonal expansion and affinity maturation (**Figure 3A, B**).^24^ Although GCs are primarily associated with B cell activity, they also harbor a distinct population of T cells, which may play a critical role in the immune response.^25^ To better understand the characteristics of these GC-associated T cells, we analyzed their TCR-Rs and compared them to those of T cells in the BCZ.

**Figure 3.**
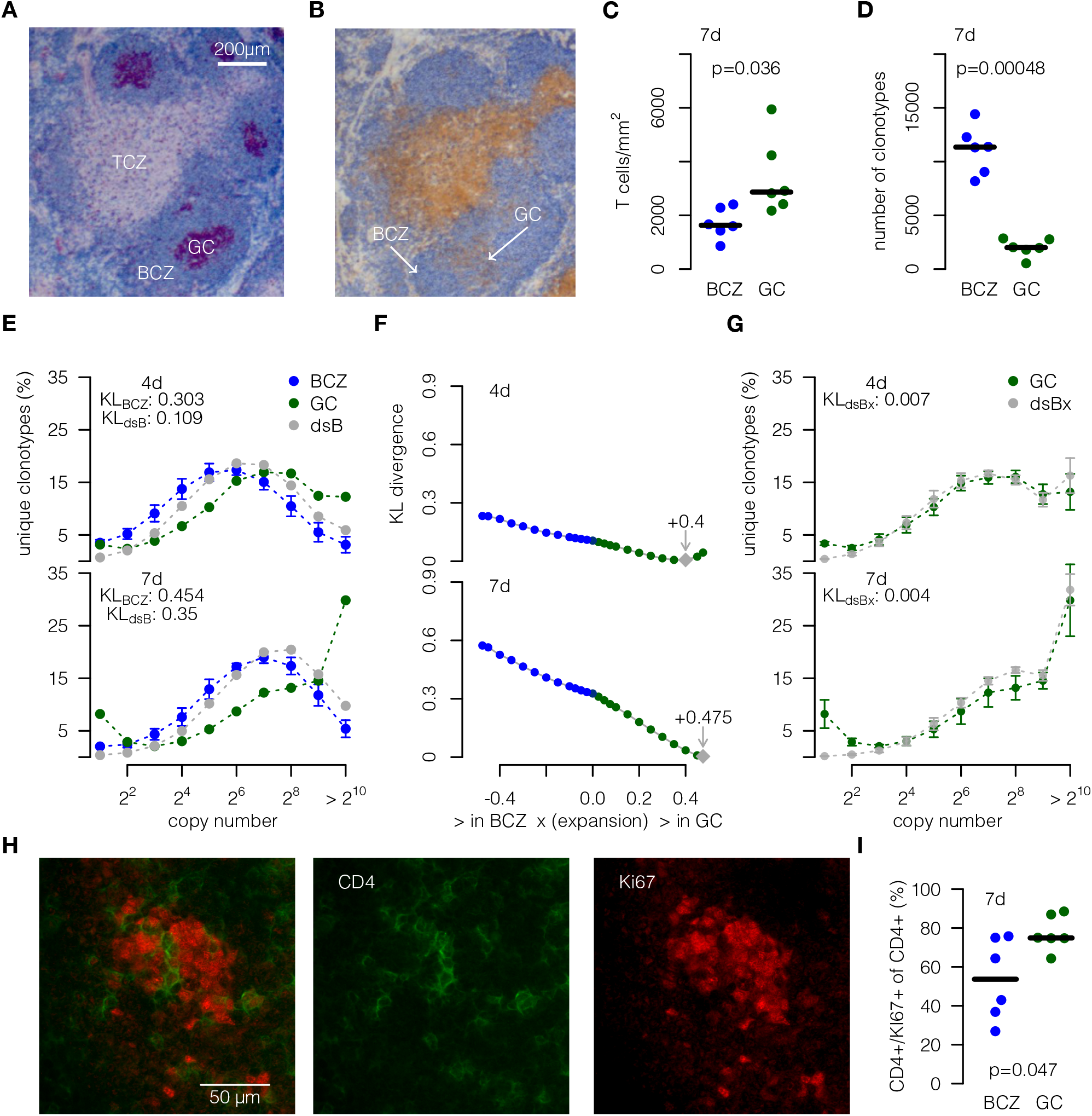
Germinal centers (GC) harbor many highly expanded T cell clones. **(A)** 7 days post-immunization (p.i.) with SRBC, splenic sections were stained for B cells (B220, blue) and proliferating cells (Ki67, red) to differentiate between the T cell zone (TCZ) and B cell zone (BCZ) and to identify GCs within the BCZ (Scale bar = 200μm). **(B)** A subsequent splenic section was stained for B cells (B220, blue) and T cells (TCRβ, brown) to quantify T cells in the BCZ and GC (arrows). **(C)** T cell densities in BCZ and GC (bars indicate means, n=6 mice, p value: two-tailed paired t test). **(D)** Number of unique amino acid CDR3β sequences per sample. Bars indicate means (n=6 mice). P value: two-tailed paired t test. **(E)** Clone size distribution at 4 days (n=7 mice) and 7 days (n=6 mice) p.i. with SRBC for BCZ, GC, and BCZ downsampled to GZ size (dsB). **(F)** Kullback-Leibler (KL) divergence for different rates of expansion, calculated on sequences with post-expansion copy number > 16. Average of 10 samples. **(G)** Clonotype size distribution for a simulated, proliferated BCZ of GC size at 4 and 7 days p.i. with SRBC. Error bars indicate standard errors (10 simulations). **(H)** Splenic sections were labeled for helper T cells (CD4, green) and proliferating cells (Ki67, red) to identify double positive cells, i.e., proliferating helper T cells in the TCZ, BCZ, and GC. A stained GC is shown, providing merged (left) and single channels (middle and right). **(I)** Percentage of proliferating helper T cells (CD4 and Ki67 positive) 7 days after immunization with SRBC (n=6). Bars indicate means. P value: two-tailed paired t test.

We found that the GC contained a significantly higher density of T cells per mm^2^ (3417 ± 1428) compared to the BCZ (1707 ± 571) 7 days p.i. (p=0.036; **Figure 3C**), in line with an earlier study of GC T cells.^25^ Despite this increased T cell density, the GC exhibited a markedly lower number of unique clonotypes than the BCZ (GC: 1993 ± 837 vs. BCZ: 11,100 ± 2247, p = 0.00048; **Figure 3D**). To determine whether these differences between the BCZ and GC repertoires were attributable solely to compartment size, we again used downsampling to match the number of unique clonotypes in the BCZ repertoire to that in the GC (dsB) at 4 and 7 days p.i. (**Figure 3E**). However, as in the previous comparison of TCZ and BCZ during the immune response, simple downsampling was insufficient to replicate the GC clone size distribution. Specifically, the GC exhibited many more clonotypes with high copy numbers than would be expected.

To further investigate the expansion dynamics, we again applied the Galton-Watson proliferation model. We tested different expansion rates to identify the best fit between the downsampled BCZ (dsB) and the GC, evaluating the similarity using KL divergence (**Figure 3F**). High expansion rates best recapitulated the GC clonotype distribution (**Figure 3F**). Specifically, a remarkably good fit between dsB and GC distributions was observed at an expansion rate of 0.4 on day 4 (KL = 0.007) and 0.475 on day 7 (KL = 0.004), suggesting high clonal proliferation within the GC (**Figure 3G**).

Further supporting this idea, immunohistochemical staining for Ki67, a proliferation marker, indeed suggested that a substantial population of the CD4^+^ T cells within the GC were proliferating (**Figure 3H**). Quantification confirmed a significantly higher proportion of Ki67^+^ CD4^+^ T cells in the GC compared to the BCZ (p = 0.047; **Figure 3I**). Together with the modeling results, these findings indicate that the distinct TCR-R clonotype size distribution in GCs is shaped at least in part by local proliferation of reactive clones rather than immigration of already expanded clones.

### No evidence for major shifts in TCR characteristics during an immune response

Thus far, our analysis has focused primarily on the distribution of clonotype sizes across the different compartments, examining how the differences between distributions can be explained by downsampling and proliferation modeling. However, in addition to these quantitative differences, we also aimed to investigate whether the sequence features (i.e., qualitative properties) of the TCR-Rs changed during the immune response.

To assess potential shifts in TCR characteristics, we analyzed the CDR3β length and P_gen_ distributions of TCR sequences in the BCZ, TCZ, and dsTx, as well as in the GC, BCZ, and dsBx. We compared these distributions at multiple time points following immunization (Figure 4A, B, Supplementary Figure S1A, B). Our analysis revealed that, despite significant changes in clonotype quantity distributions, the CDR3β length and P_gen_ profiles remained largely stable throughout the immune response and across compartments, also when downsampling and modeling proliferation. These findings indicate that while T cell expansion dynamics shift substantially during an immune response, leading to changes in clonotype abundance, the sequence features of the TCR-Rs remain largely unchanged between compartments.

**Figure 4.**
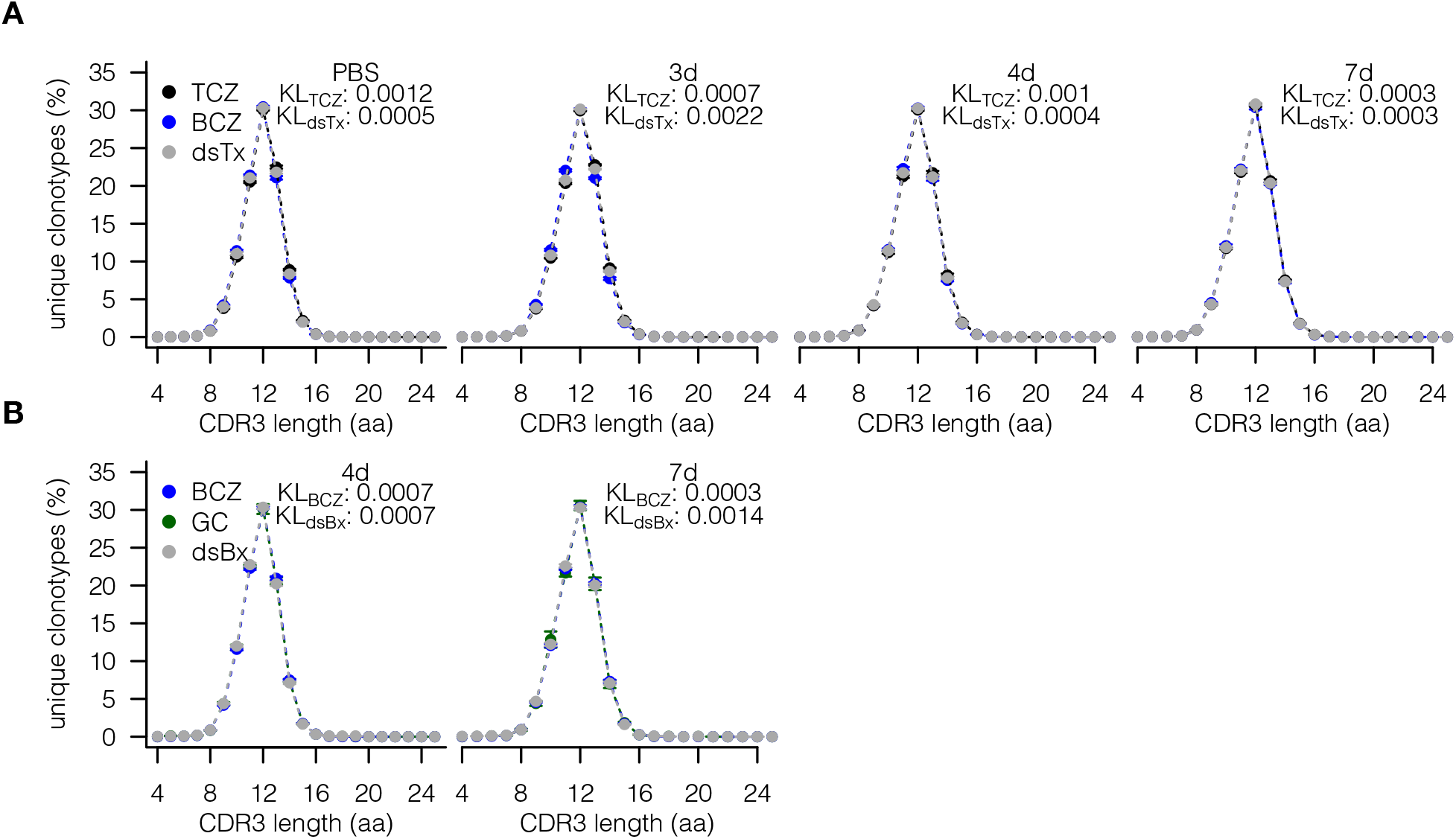
CDR3β length distribution remains stable despite differing clone size distributions across compartments and timepoints. **(A)** CDR3β length distribution in the T cell zone (TCZ), B cell zone (BCZ), and down-sampled, proliferated TCZ (dsTx) for PBS (n=8 mice), 3d (n=6 mice), 4d (n=7 mice), and 7d (n=6 mice) after immunization with SRBC. **(B)** CDR3β length distribution in BCZ, GC and down-sampled, proliferated BCZ (dsBx) for 4d, and 7d after immunization with SRBC.

### The GC TCR-R contains clusters of similar sequences that can be traced back to the early immune response

After examining overall clonotype distributions and sequence features, we next focused on the composition of individual TCR sequences within the GC. Specifically, since TCRs responding to the same antigen often have similar sequences,^26, 27^ we asked whether the GC harbors clusters of similar TCR sequences across mice and whether these were already present earlier in the immune response in other compartments.

To identify related TCRs, we pooled all TCR sequences from mice at day 7 p.i. and performed a clustering analysis based on sequence similarity. Sequences were connected if they shared the same V gene and J gene and their CDR3β junction sequence differed in at most one amino acid. This approach revealed distinct clusters of similar sequences within the GC (**Figure 5A**). Most clusters contained sequences from multiple mice, suggesting that these TCRs were responding to common antigenic stimuli (**Figure 5B**).^26,27^ Three of the four largest clusters containing sequences from multiple mice shared the same V gene (TRBV3) and a J gene from the same J gene locus (TRBJ2). However, we also observed clusters in the GC that only occurred in one mouse, indicative of a more individualized (private) immune response to at least some epitopes of the complex SRBC antigen.^17,18,23,28^

**Figure 5.**
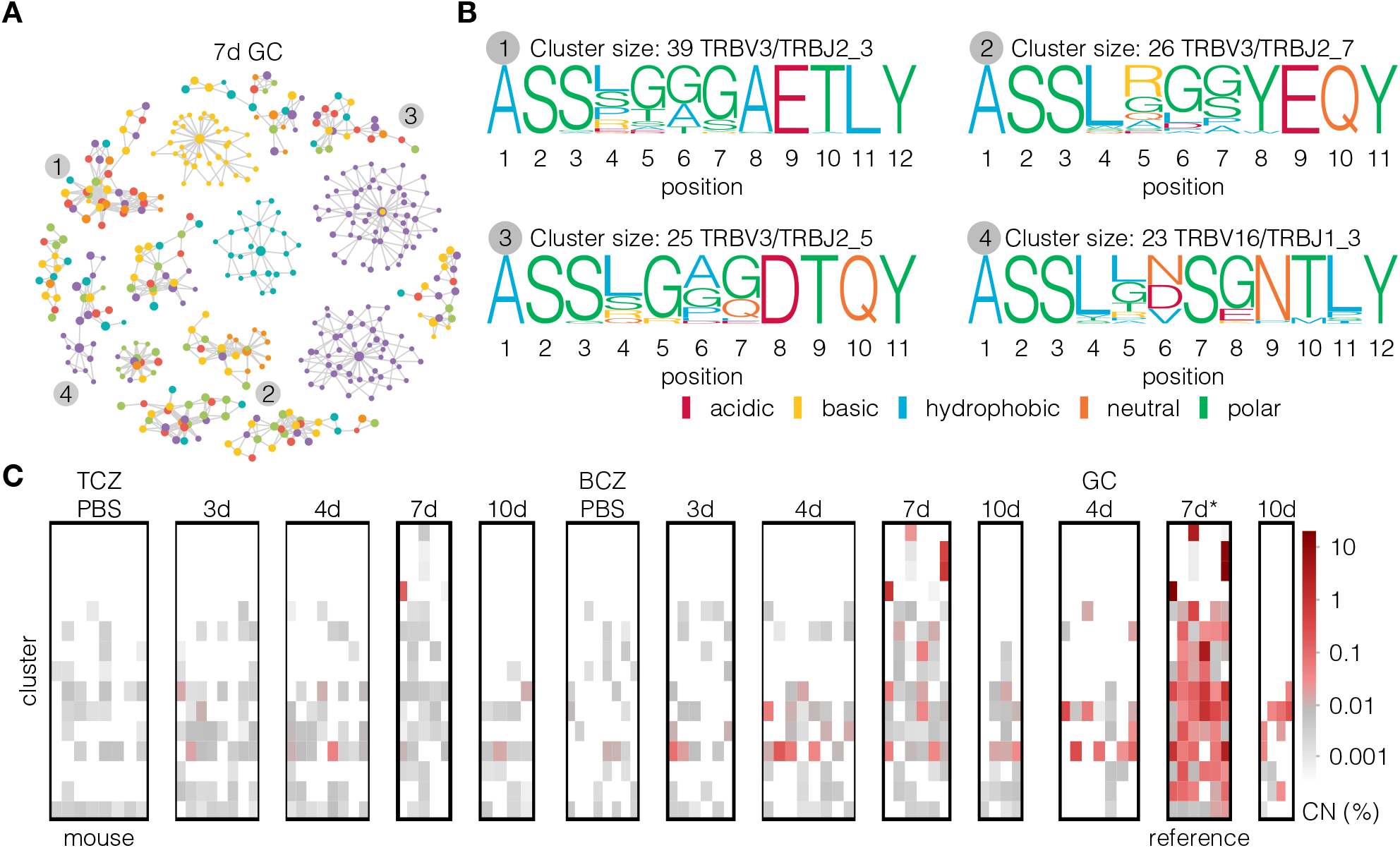
The Germinal Center (GC) harbors many similar CDR3β sequences that likely respond to the same antigens and can be traced back to the early immune response. **(A)** Clusters of T cells in GC mice 7 days post-immunization (n=6 mice). TCRs are linked if their Hamming distance is one or less. Clusters of 15 or more distinct sequences are shown. Colors represent different mice. **(B)** Sequence logos for the four largest clusters containing sequences from multiple mice. **(C)** Percentage of total copy numbers associated with sequences in each GC 7d cluster per mouse. Sequences are tracked through the compartments based on their CDR3β nucleotide sequence, V gene, and J gene. The color scale is logarithmic, with 0.0001 added to each copy number percentage.

To trace these probably antigen-reactive sequences back to earlier timepoints in the immune response, we searched for the sequences from the GC clusters across all samples and time points. By mapping CDR3β nucleotide sequences, V genes, and J genes, we found that a substantial proportion of the clonotypes from these GC clusters could be detected in earlier immune response stages within the BCZ and TCZ. The copy numbers of the clusters increased over time, indicating rapid expansion during the immune response, primarily within the GC (**Figure 5C**). These findings suggest that the GC TCR-R is shaped by the selective expansion of specific T cell clones that likely emerged early in the immune response in the TCZ and BCZ. The presence of highly similar TCR clusters in multiple mice suggests that these clones respond to shared antigenic targets.

## Discussion

The spleen is a key SLO that plays a central role in initiating and regulating immune responses, particularly to blood-borne antigens with T cells playing a crucial role.^29^ Due to the high number of unique TCRs and their variability, analyses of the TCR-Rs are challenging, and sensitive tools are needed to monitor changes within these highly diverse TCR-Rs. Comparing TCR-Rs between compartments of the spleen brings an extra challenge due to the varying densities of T cells in the different compartments. In this study, we used down-sampling and proliferation modeling to quantify the differences between the TCZ, BCZ, and GC under naïve conditions and following an immune response to SRBC. Our results indicate that observed differences in TCR-Rs across compartments can be almost entirely attributed to variations in compartment size and proliferation dynamics.

First, we compared the TCR-Rs between TCZ and BCZ compartments under both naïve conditions and during an SRBC-induced immune response. In the absence of antigenic stimulation, both the T cell density and clonotype diversity in the BCZ were significantly lower than those in the TCZ. However, after adjusting for compartment size through downsampling, the TCR-Rs in both compartments became highly similar. This indicates that the BCZ does not harbor a unique repertoire but instead represents a reduced subset of the larger TCR pool present in the TCZ that may have migrated to the BCZ randomly.^19^ After antigenic stimulation, the repertoires of BCZ and TCZ diverged in ways that down-sampling alone could not explain. Using a proliferation model based on the Galton-Watson process, we determined that the differences observed 3 and 4 days p.i. were explainable by differing proliferation rates between the compartments: Initially, the TCZ exhibited higher proliferation, but by day 4, the BCZ showed a larger number of expanded clonotypes. This could either mean that activated T cells migrated into the BCZ and expanded there, or that expanded T cells from the TCZ migrated into the BCZ. By day 7, these differences between the compartments has largely disappeared. Our findings align with previous observations of transient TCR compartmentalization following antigenic challenge. In particular, the shifting-mosaic model proposed in prior work suggests that T cell migration normally dominates, but proliferation can briefly outpace it upon immune activation.^19,30^ Our proliferation modeling further supports this idea.

Second, we explored the GC compartment, a critical site for B cell expansion and maturation during an immune response. Despite increased T cell density in the GC compared to the TCZ, its TCR diversity was significantly lower than in the BCZ, indicating selective recruitment, local expansion, or both of antigen-reactive T cells. Clustering analysis identified groups of highly similar TCR sequences within the GC across different mice.^26,27,31^ Our proliferation modeling demonstrated that high expansion rates best explained the observed clonotype distribution compared to the BCZ, underscoring the role of intense clonal proliferation in shaping the GC-resident TCR-R. Immunohistochemistry staining for Ki67 further supported this, showing a high density of proliferative CD4^+^ T cells in the GC.^9^

Finally, we examined the qualitative nature of the TCR-Rs. Despite substantial shifts in clonotype abundance across compartments, the sequence characteristics of the TCR-Rs, such as CDR3β length length, remained stable. Therefore, despite highly dynamic T cell expansion and migration, the qualitative properties of the TCR-Rs as a whole appear to have been preserved.

One limitation of our approach is that the Galton-Watson model is relatively simplistic, as it only accounts for growth through stochastic proliferation. Consequently, it does not capture the emergence of new, low-copy number cells, making it difficult to accurately model this part of the TCR-R. Additionally, our model does not distinguish whether T cells initially proliferated in one compartment and later migrated to another or if proliferation occurred exclusively within a single compartment, limiting our ability to resolve the exact sequence of these events. Despite these limitations, the basic principles of proliferation captured by the model led to a remarkable fit to the observed clonotype sizes in TCZ and GC. Future studies could refine this approach by incorporating, for instance, cell death and the introduction of new cells in later steps of the branching process,^32,33^or agent-based simulations,^34^ focussing on the behavior of single cells and the sequence of events. These improvements would allow for a better capture of additional factors concerning selection, migration, and proliferation. Further, single-cell data could be used to resolve the TCRα chain and get more complete information on clonotype identity.

Overall, our study highlights despite the apparent major differences between the TCR-Rs of splenic compartments, there is in fact a remarkable consistency between them. This consistency may only become apparent once the effects of compartment size and proliferation are explicitly factored into the analysis.

## Acknowledgements

This research was supported by an NWO Vidi grant (VI.Vidi.192.084) to JT. JW was funded by the German Research Foundation (DFG) within the framework of the TR-SFB654 project C4 at the University of Lübeck. We want to thank P. Lau for performing the light microscopy.

## Author contributions

In collaboration with CT, FB conducted the TCR-R data analyses under the supervision of JT and JW, with key input from AS and LKS. CT, AS and LKS conducted the mouse experiments. RP carried out the NGS analyses. LKS performed the light microscopy and laser-microdissection. AS was responsible for cell staining, imaging, and analyses. The manuscript was written by FB, CT, AS, and JT with feedback from all other authors. All authors read and approved the final manuscript.

## Data and code availability

The processed TCR sequences and the code required to reproduce the analyses reported in this manuscript are available on Zenodo.^35^

## Supplementary Figures

**Supplementary Figure S1.**
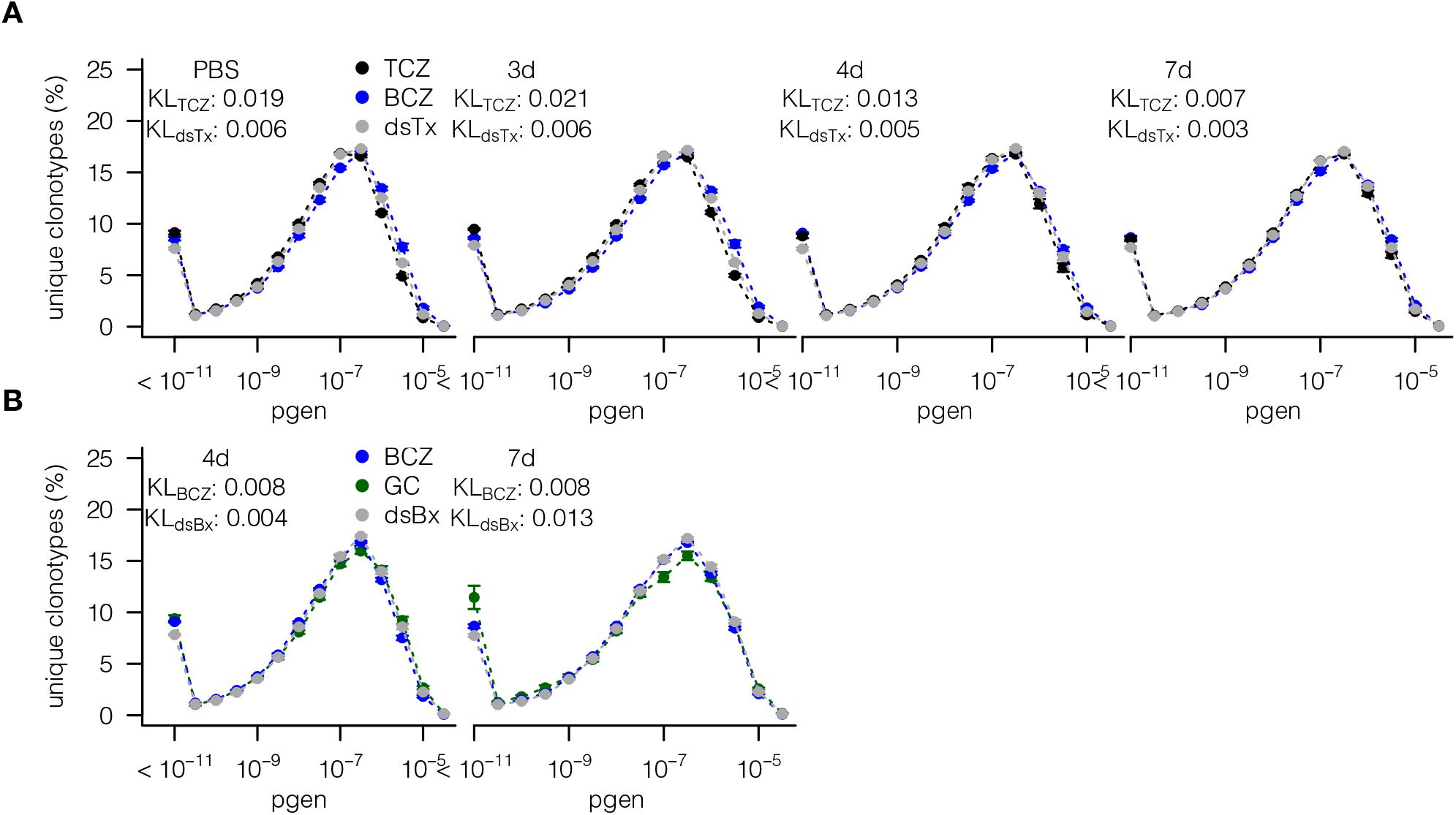
CDR3β pgen distribution remains stable despite differing clone size distributions across compartments and timepoints. **(A)** CDR3β pgen distribution in the T cell zone (TCZ), B cell zone (BCZ), and down-sampled, proliferated TCZ (dsTx) for PBS (n=8 mice), 3d (n=6 mice), 4d (n=7 mice), and 7d (n=6 mice) after immunization with SRBC. **(B)** CDR3β pgen distribution in BCZ, GC and down-sampled, proliferated BCZ (dsBx) for 4d, and 7d after immunization with SRBC.

